# Regulatory T and activated T cells in the chronic unpredictable mild stress mice: sex differences and effects of treatments

**DOI:** 10.1101/2025.02.04.636379

**Authors:** Mengqi Niu, Yingqian Zhang, Abbas F. Almulla, Tangcong Chen, Yueyang Luo, Mengdie Li, Juan Li, Yaoyin Zhang, Yulan Huang, Michael Maes

## Abstract

**Background:** Major depressive disorder (MDD) is associated with immune dysregulation, including T-cell activation and reduced regulatory T cells. Pharmacological treatments like fluoxetine, simvastatin, curcumin, and S-adenosylmethionine (SAMe) show promise in alleviating depressive symptoms, but their effects on immune responses are not fully understood.

**Methods:** This study explored immune status in MDD using the chronic unpredictable mild stress (CUMS) mouse model. It examined how fluoxetine, simvastatin, curcumin, and SAMe affected immune responses and depressive behaviors. The study also investigated sex differences. Six groups, each with 12 mice (6 males and 6 females), were included: control, CUMS, and drug treatments. Male and female mice were housed separately in sex-specific cages throughout the experiment, with three mice per cage. Behavioral tests included sucrose preference, forced swimming, open field, and active avoidance. Immune cell markers were analyzed by flow cytometry.

**Results:** CUMS mice showed immune imbalance, with Treg depletion (*p* = 0.047) and a higher CD154/CD152 ratio (*p* = 0.006). All treatments improved depressive-like behaviors (all *p* < 0.05), with SAMe normalizing the CD154/CD152 ratio (*p* = 0.009). Furthermore, female mice exhibited lower levels of Treg-related immune regulation and T-cell activation compared to males (both *p* < 0.001), along with a higher CD154/CD152 ratio (*p* = 0.008).

**Conclusion:** This study highlights that curcumin, simvastatin, fluoxetine, and SAMe alleviate depressive-like behaviors, while SAMe also reduces the CD154/CD152 ratio. These findings reveal marked sex differences in T-cell activation and immune regulation, highlighting the importance of considering sex as a biological variable in CUMS research.

## 1. Introduction

Major Depressive Disorder (MDD) is a common psychiatric condition, often associated with immune system dysregulation, which includes excessive T-cell activation (elevated CD69+, CD71+, CD40L+/CD154+, HLA-DR+) and a reduction in regulatory T cells (Tregs, CD4+CD25+FoxP3+), disrupting immune homeostasis ^1–5^.

In MDD, the upregulation of CD154 (a T-cell activation marker) enhances the production of pro-inflammatory cytokines (IL-6, TNF-α) ^1–5^, while the downregulation of CD152 (CTLA-4, an inhibitory receptor) impairs immunoregulation and immune tolerance ^6^. The consequent immune imbalance is observed both in patients and in chronic unpredictable mild stress (CUMS) models ^3,7–10^. The CUMS model, developed by Willner et al., exposes mice to unpredictable mild stressors to mimic human chronic stress, and can induce multidimensional behavioral alterations, including anhedonia- like behavior, increased forced-swim immobility, anxiety-like behavior, and cognitive impairment; it is widely used to study immune dysregulation and treatment responses in depression ^11–13^. Accordingly, the sucrose preference test (SPT), forced swimming test (FST), open field test (OFT), and active avoidance test (AAT) were selected to assess anhedonia-like behavior, behavioral despair, locomotor/exploratory and anxiety-like behavior, and conditioned avoidance behavior, respectively ^14,15^.

Recent evidence conceptualizes MDD as involving interconnected neuroimmune, metabolic, and oxidative stress (NIMETOX) pathways, including T-cell dysregulation, lipid-metabolic abnormalities, oxidative/nitrosative stress, and mitochondrial dysfunction ^16^. These interacting abnormalities provide a mechanistic rationale for pharmacological interventions targeting complementary components of the NIMETOX network ^16,17^. Fluoxetine, a selective serotonin reuptake inhibitor (SSRI), not only regulates serotonin levels but also reduces pro-inflammatory cytokine levels and enhances anti-inflammatory responses, contributing to its therapeutic potential ^18,19^. Research has shown that simvastatin modulates the Th17/Treg balance ^20^. Beyond its lipid-lowering action, simvastatin has also shown antidepressant-like and immunomodulatory effects in chronic-stress models, involving restoration of hippocampal NMDAR-related synaptic plasticity and inhibition of P2X7/NLRP3 inflammasome signaling ^21,22^. Curcumin, a polyphenol with antioxidant and anti-inflammatory properties, inhibits T-cell activation, reduces Th1/Th17 differentiation, and promotes Treg generation ^23^. It has also shown antidepressant-like effects in preclinical models, associated with attenuation of inflammatory responses through PI3K/Akt signaling and modulation of neuroinflammation-related MAPK signaling ^24,25^. S-adenosylmethionine (SAMe), an endogenous metabolite, has the ability to restore neurotransmitter activity (serotonin (5-HT) and dopamine (DA)) and membrane fluidity ^26^. Its efficacy is comparable to that of traditional antidepressants, making it a potential alternative treatment for depression ^27^. Accordingly, the four agents were selected as complementary pharmacological interventions within the NIMETOX framework: fluoxetine as an established antidepressant reference, simvastatin as a clinically used repurposing drug with immunometabolic actions, curcumin as a natural antioxidant and immunomodulatory compound, and SAMe as an endogenous metabolic adjunct with antidepressant potential.

Sex differences in MDD are significant, with women being 2-3 times more likely to develop the disorder than men, likely due to differences in immune regulation. Studies have shown that women have fewer Tregs, an increase in Th1 and CD8+ T cells ^28^, and exhibit stronger pro-inflammatory responses (e.g., elevated TNF-α/IL-10 and iNOS/Arg-1 ratios) under CUMS ^29^. These differences may be driven by estrogen’s immunostimulatory effects and testosterone’s immunosuppressive effects ^30^. These disparities, along with genetic and gut microbiota factors, may partially explain the higher prevalence and more severe symptoms of MDD in women ^31^. Although sex differences, drug effects, and T-cell responses have all been studied previously in CUMS models, the specific association between sex and immune responses, particularly the balance between activated T cells and Treg-mediated immune regulation, has not been systematically examined in CUMS mice.

This study aimed to investigate the role of T-cell activation and Treg depletion in depression using the CUMS mouse model, with a focus on how fluoxetine, simvastatin, curcumin, and SAMe modulate immune responses and alleviate depressive symptoms. Furthermore, we explored the impact of sex differences on immune responses and drug efficacy, providing theoretical insights for developing sex-specific therapeutic strategies for MDD.

## 2. Materials and methods

### 2.1 Animals and chronic unpredictable mild stress

Seventy-two 3-week-old C57BL/6N mice (half male, half female; body weight at arrival, 7.0–12.6 g) were obtained from GemPharmatech (Chengdu, China) and housed in standard conditions (12-hour light/dark cycle, 23 ± 1°C) with standard chow and water. All animals were housed in sex-specific cages throughout the study, with three mice per cage. The mice were assigned to six groups (n = 12 per group; 6 males and 6 females): control, CUMS, CUMS plus fluoxetine, CUMS plus simvastatin, CUMS plus curcumin, and CUMS plus SAMe. Each group therefore comprised four cages: two male cages and two female cages. The experimental design is shown in **Fig. 1**. All procedures were approved by the Institutional Animal Care and Use Committee (IACUC) of Sichuan Junhui Biotech Co., Ltd. (approval number: IACUC-202411-4-001).

**Fig. 1.**
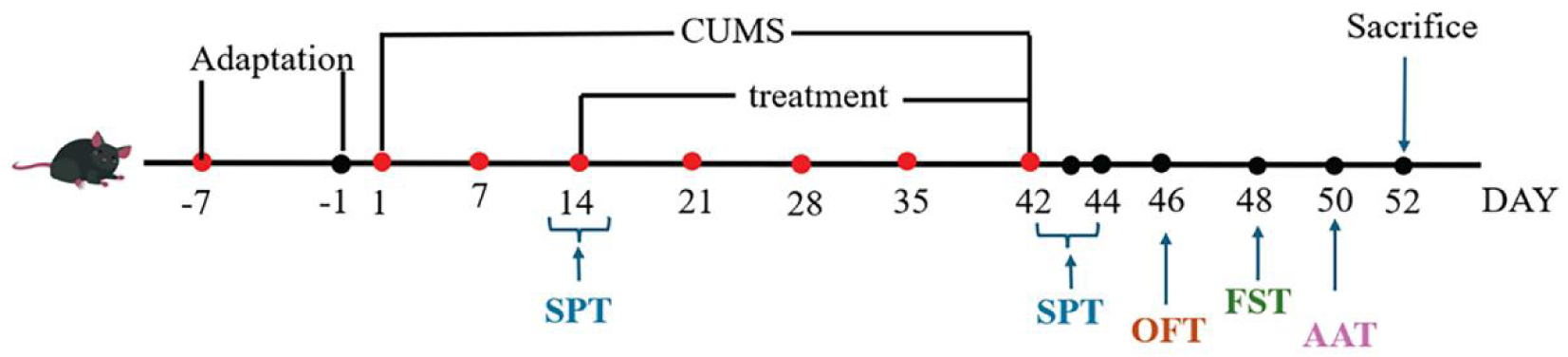
The experimental design. Schematic diagram of the experimental design. After a 7-day adaptation period in the animal facility (from day -7 to -1), the mice were randomly assigned to the control group (12 mice, half male and half female) and the CUMS group (60 mice, half male and half female) for model induction. After two weeks of CUMS modeling, SPT was performed, and the mice were then randomly divided into five groups (CUMS, Fluoxetine, Simvastatin, Curcumin, and SAMe, with 12 mice per group, half male and half female). Mice continued receiving CUMS modeling and treatment for 4 weeks (from day 14 to day 42), followed by behavioral testing (SPT on day 42, OFT on day 46, FST on day 48, AAT on day 50). Euthanasia was performed on day 52. One male mouse in the CUMS plus SAMe group died during CUMS modeling, whereas no other deaths occurred.

For CUMS modeling, mice were exposed to two daily stressors, including tilted cages (45°), noise and light flashes, wet bedding, social isolation, water deprivation, and restraint. During the 6-week CUMS modeling period, animals were exposed to two distinct mild stressors per day, with stressor sequences randomized weekly and no stressor repeated on consecutive days to minimize habituation (see **Table 1**) ^32^.

**Table 1.** Chronic unpredictable mild stress (CUMS) stressors and exposure parameters.

| Stressor domain | Stressor | Key parameters | Exposure duration | Schedule constraints | Source |
| --- | --- | --- | --- | --- | --- |
| Sensory | Auditory/visual stimulation | 85 dB white noise + 3 Hz flashing light | 1 h | Weekly randomization; no consecutive-day repetition | Li et al., 2022<sup>32</sup> |
| Psychosocial | Social isolation | Single housing | 24 h | Weekly randomization; no consecutive-day repetition | Li et al., 2022<sup>32</sup> |
| Environmental | Cage tilt | 45° tilt | 24 h | Weekly randomization; no consecutive-day repetition | Li et al., 2022<sup>32</sup> |
| Deprivation | Water deprivation | Water removed | 24 h | Weekly randomization; no consecutive-day repetition | Li et al., 2022<sup>32</sup> |
| Environmental | Wet bedding | 200 mL of water added to the bedding | 24 h | Weekly randomization; no consecutive-day repetition | Li et al., 2022<sup>32</sup> |
| Physical | Restraint | Ventilated 50 mL tube | 2 h | Weekly randomization; no consecutive-day repetition | Li et al., 2022<sup>32</sup> |

Animals in the CUMS condition were exposed to two distinct mild stressors per day for 6 consecutive weeks. Stressors were randomized weekly, and the same stressor was not applied on consecutive days to minimize habituation. Control animals were housed under identical conditions without stress exposure.
Abbreviations: CUMS, chronic unpredictable mild stress.

### 2.2 Treatment

Fluoxetine, Simvastatin, Curcumin, and SAMe were administered for 28 days after 2-week CUMS (**Fig. 1**). Fluoxetine (Aladdin, CAS No.: 54910-89-3): 20 mg/kg, orally, dissolved in normal saline ^32^. Simvastatin (Aladdin, CAS No.: 79902-63-9): 10 mg/kg, orally, dissolved in 1% DMSO (Sigma, CAS No.: 67-68-5)^33^. Curcumin (Aladdin, CAS No.: 458-37-7): 20 mg/kg, intraperitoneally, dissolved in peanut oil ^34^. SAMe (Solarbio, CAS No.: 97540-22-2): 100 mg/kg, orally, dissolved in normal saline ^35^. More detailed information regarding compound preparation, dose, solvent/vehicle, route of administration, and treatment schedule can be seen in **ESF Table 1**.

### 2.3 Sucrose preference test (SPT), forced swimming test (FST), open field test (OFT) and active avoidance test (AAT)

Behavioral testing and scoring were performed by experimenters blinded to group assignment. The timing and order of the behavioral tests are shown in **Fig. 1**.

SPT: A 1% sucrose solution and distilled water were provided in two separate bottles in each cage for two consecutive days. After 6 hours of food and water deprivation, the mice were housed individually on the third day. One bottle contained 1% sucrose solution, while the other contained distilled water. The positions of the sucrose and water bottles were counterbalanced to minimize side preference. The bottles were weighed before placement, and after 12 hours, the remaining liquid was weighed. The sucrose preference percentage was calculated using the formula: Sucrose Preference (%) = (Sucrose Consumption / Total Fluid Consumption) × 100%.

FST: The mice were placed in a transparent cylindrical container (diameter: 10 cm, height: 30 cm) filled with water (depth: 20 cm, temperature: 25 ± 1°C), so that they could not touch the bottom. Data were recorded for 6 minutes, with the last 4 minutes used for analysis. Immobility was defined as making only the minimal movements necessary to keep the head above water.

OFT: Each mouse was gently placed in the center of an open field box (50 × 50 × 50 cm) under approximately 50 lux illumination for 10 min to allow it to explore freely. When setting the open field area, we divided it into 4 × 4 grids, and the middle four grids (25 × 25 cm) were defined as the center area. Movement was recorded and analyzed using an automated tracking system (Cleversys, USA). Total distance travelled, the duration of center-zone bouts, and the duration spent in the corner zones were recorded as distance travelled (m), duration in center (seconds), and duration in corner (seconds), respectively.

AAT: The test was conducted in an apparatus (30 × 30 × 20 cm) consisting of equally sized light and dark compartments connected by an opening (3 cm high × 4 cm wide). The light compartment was illuminated at 200–400 lux, and animal behavior was recorded using infrared cameras. Mice were acclimated to the behavioral testing room for 24 h before testing. During both the training and testing phases, mice were initially placed in the dark compartment. During training, mice received foot shocks while in the dark box. Foot shocks were delivered at 0.2–0.8 mA (100 V, 50–60 Hz AC) for 1– 2 s, using the minimum intensity required to elicit withdrawal and vocalization. Upon escaping to the lightbox to avoid the shock, the mice were immediately removed and returned to their cage. Two hours later, during the test phase, the mice were reintroduced to the dark box without foot shocks. The duration spent in the light box during the 5-min test was recorded using infrared cameras and analyzed using TopScan software (Cleversys, USA). Both compartments were cleaned with alcohol between animals.

### 2.4 Spleen sampling and flow cytometry

Mice were placed in a closed chamber, and CO₂ was gradually introduced at a displacement rate of 15–20% of the chamber volume per minute to induce deep anesthesia. Adequate anesthesia was confirmed by the loss of the righting reflex and pedal withdrawal reflex. While the mice remained under deep anesthesia, spleens from eight randomly selected mice per group (4 males and 4 females) were collected for flow-cytometric analysis. After sample collection, euthanasia was completed by cervical dislocation, and death was confirmed by the absence of spontaneous respiration and reflex responses.

The spleens were mechanically prepared into single-cell suspensions at a density of 1 × 10⁶ cells/mL. A 100 μL aliquot of the cell suspension was placed in a flow cytometry tube for further analysis. Two standardized flow cytometric panels were used for immune phenotyping: a seven-color surface panel (CD4-eFluor 450, CD8-BV650, CD40-PE-Cy5, CD69-BV785, CD71-PE, HLA-DR-FITC, CD154-APC) and a five-color surface/intracellular panel (CD4-eFluor 450, CD25-Alexa Fluor 488, GARP-Super Bright 702, CD152-APC, Foxp3-PE), with all antibodies purchased from their corresponding commercial vendors. For surface staining, the cell suspensions were incubated with the corresponding antibody cocktails at 4°C for 30 min and then washed with PBS. For intracellular Foxp3 staining, cells were fixed and permeabilized for 40 min, washed with 1× permeabilization buffer, and incubated with Foxp3-PE at 4°C for 30 min before acquisition.

Sample acquisition was conducted on a ZE5 flow cytometer (Bio-Rad, USA), and data were analyzed using Everest v3.1.18.0 software (Bio-Rad, USA). The detailed gating strategy is shown in **Fig. 2**. All analyses were performed in accordance with established flow cytometry guidelines ^36^.

**Fig. 2.**
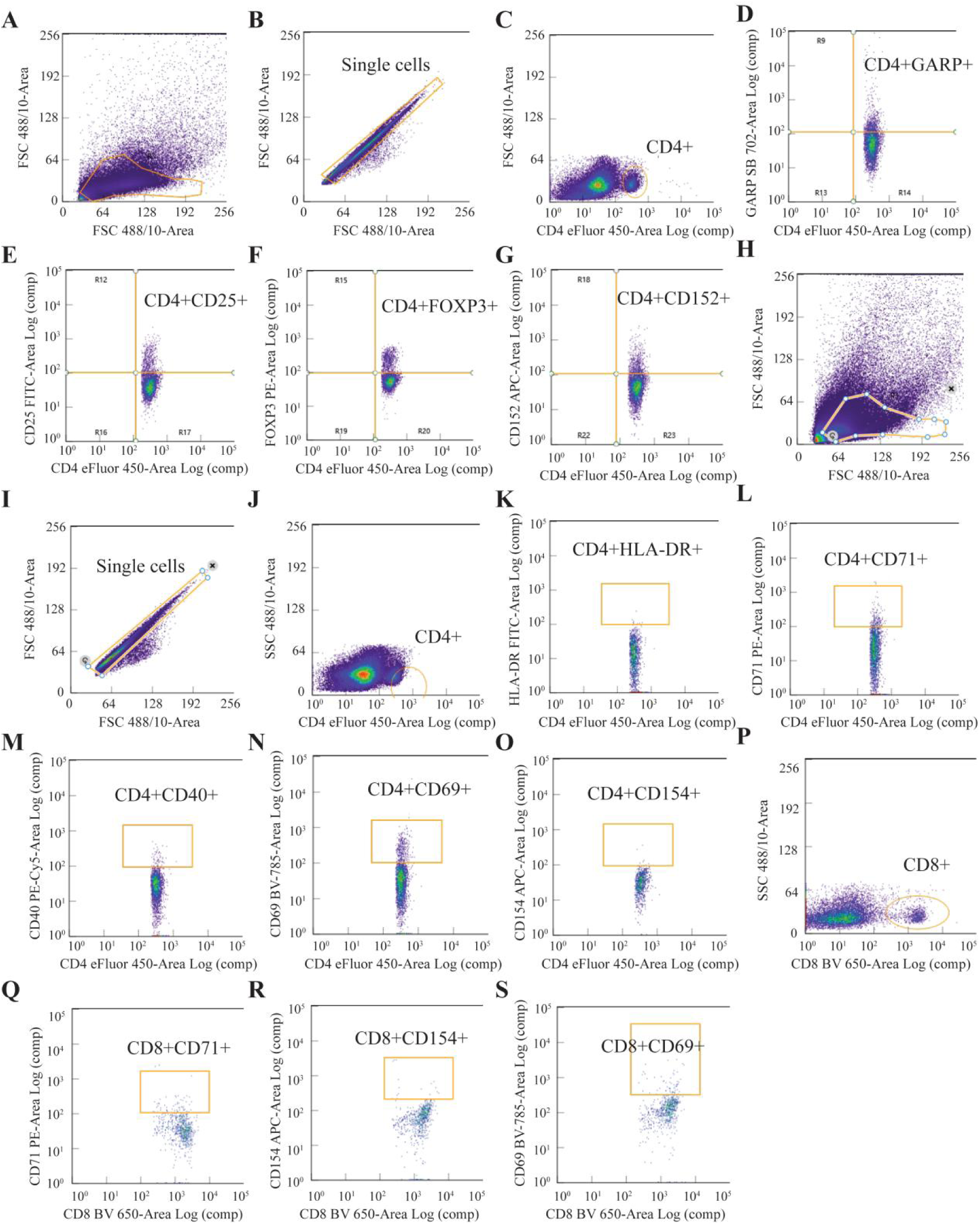
Flow cytometry gating strategy. This figure illustrates the gating strategy used in flow cytometry to identify specific immune cell populations. The gates represent sequential steps for selecting regulatory CD4+ T cells and their subsets (A-G), activated CD4+ T cells and their subsets (H-O), and CD8+ cells and subsets (H-I, P-S). Spleen samples from eight mice per group (4 males and 4 females) were analyzed by flow cytometry.

### 2.5 Statistical analyses

All analyses were conducted utilizing univariate general linear model analyses with six groups (control, CUMS, Fluoxetine, Simvastatin, Curcumin, SAMe) and sex as fixed factor. We also incorporated the treatment by sex effect. We employed protected least significant difference tests to analyze pairwise comparisons, assessing the differences between the control and CUMS conditions, as well as among the CUMS conditions and the effects of four treatments. Should the latter effects prove significant, we also analyzed the differences between the treatment and control conditions. Simple-effects analyses were performed only when the treatment group × sex interaction was statistically significant. The data were standardized (z-transformed) and z unit-based composite scores were computed. The statistical analysis for this study was performed using version 28 of IBM SPSS for Windows. All analyses were conducted using two-tailed testing. A p-value of 0.05 or lower was deemed indicative of statistical significance.

## 3. Results

In this study, we utilized a factorial general linear model (GLM) to analyze the relationships between treatment groups (control, CUMS, Fluoxetine, Simvastatin, Curcumin, SAMe) and sex (male and female). The factorial GLM combines the concepts of GLM and factorial design. This model allows us to analyze both the individual effects of each factor and their interactions, i.e., how combinations of factors jointly influence the outcome. Specifically, we conducted the analysis at three levels: First, we examined the differences among the treatment groups. The results confirmed the successful establishment of the CUMS model. This was indicated by significant differences in SPT (*p* < 0.001), FST (*p* = 0.001), OFT (*p* < 0.01) and AAT (*p* = 0.023) between the CUMS and control condition.

Furthermore, the factorial GLM analysis revealed significant differences between the CUMS group and the treatment groups in behavioral and immune measures. Second, we considered the impact of sex, and observed sex-related differences in immune cell subsets. Finally, the analysis showed that there were no significant interactions between sex and treatment groups in any of the behavioral outcomes or immune cell subsets.

### 3.1 Behavioral Differences Between the Control, CUMS, Fluoxetine, Simvastatin, Curcumin, and SAMe Groups

We used a variety of behavioral tests to evaluate depression-like behaviors in CUMS model. Specifically, SPT assessed anhedonia (**Fig. 3A**); FST evaluated behavioral despair (**Fig. 3B**); OFT measured autonomous activity ability and anxiety-like behaviors (**Fig. 3C–3E**), and AAT assessed learning and memory capabilities (**Fig. 3F**). The SPT conducted on day 14, before treatment initiation, showed that each of the five CUMS-exposed groups had significantly lower sucrose preference than the control group (all *p* < 0.01). At the experimental endpoint, mice in the CUMS group exhibited the lowest sucrose preference (**Fig. 3A**), the longest immobility time (**Fig. 3B**), the shortest distance traveled, the least time spent in the center, and the longest time spent in the corners in the OFT (**Fig. 3C–3E**), as well as the shortest duration in the lightbox in the AAT (**Fig. 3F**). These findings confirm the successful establishment of the CUMS model (all *p* < 0.05).

**Fig. 3.**
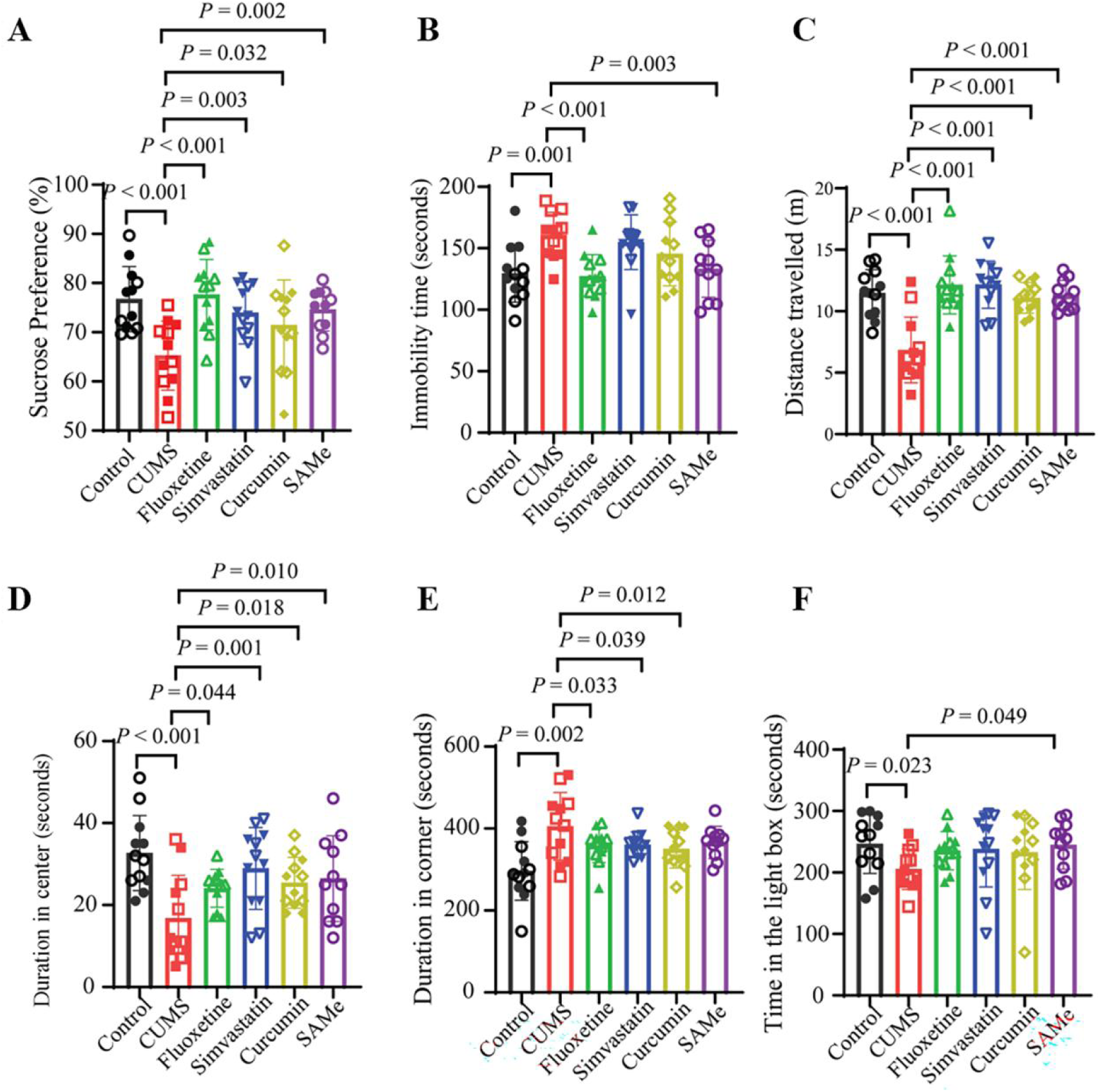
Behavioral differences among the control, CUMS, fluoxetine, simvastatin, curcumin, and SAMe groups. **(A)** Sucrose preference in the SPT. **(B)** Immobility time in the FST. **(C-E)** Differences in OFT among six groups of mice: control, CUMS, fluoxetine, simvastatin, curcumin, and SAMe. **(F)** Duration spent in the light box in the AAT. Each symbol represents one mouse as a biological replicate; solid and hollow symbols represent male and female mice, respectively. For the endpoint behavioral analyses shown in A–F, n = 12 per group (6 males and 6 females) for the control, CUMS, fluoxetine, simvastatin, and curcumin groups, and n = 11 (5 males and 6 females) for the SAMe group. Data are presented as mean ± standard error of the mean (SEM). Each behavioral outcome was analyzed using a factorial general linear model (GLM) with treatment group and sex as fixed factors. When a significant treatment-group effect was detected, protected least significant difference (LSD) tests were used for pairwise comparisons. All tests were two-tailed, and exact *P* values are shown above the corresponding comparison brackets.

Following treatment, sucrose preference significantly increased in the Fluoxetine, Simvastatin, Curcumin, and SAMe groups compared to the CUMS group (**Fig. 3A**, all *p* < 0.05). In the FST, immobility time was significantly reduced in both the Fluoxetine and SAMe groups (**Fig. 3B**, all *p* < 0.01). In the OFT, all four treatments significantly increased distance traveled and time spent in the center (**Fig. 3C–3D**, all *p* < 0.05), while Fluoxetine, Simvastatin, and Curcumin significantly reduced time spent in the corners (**Fig. 3E**, all *p* < 0.05). In the AAT, SAMe markedly increased the duration spent in the lightbox (**Fig. 3F**, *p* < 0.05). In summary, the four treatments alleviated CUMS-induced anhedonia, reduced autonomous activity ability, and anxiety-like behaviors to varying degrees. Notably, Fluoxetine and SAMe also alleviated behavioral despair, while SAMe further improved learning and memory impairments induced by CUMS.

### 3.2 Immune Cell Differences Between the Control, CUMS, Fluoxetine, Simvastatin, Curcumin, and SAMe Groups

Differences in immune cell markers among the Control, CUMS, Fluoxetine, Simvastatin, Curcumin, and SAMe groups are illustrated in **Fig. 4**. **Fig. 4A** displays the differences in composite indices across groups. The indices include z(CD154-CD152) or the ratio of CD154 versus CD152, which is calculated as z(CD4+CD154+) + z(CD8+CD154+) − z(CD4+CD152+); z(TACT), which represents T-cell activation, calculated as z(CD4+HLA-DR+) + z(CD4+CD71+) + z(CD4+CD69+) + z(CD4+CD40+) + z(CD4+CD154+); and z(Treg), which represents regulatory T cells, calculated as z(CD4+FOXP3+GARP+) + z(CD4+CD25+FOXP3+GARP+) + z(CD4+CD25+FOXP3+CD152+). Furthermore, the ratio of activated T versus Treg cells or z(TACT-Treg) is computed as Z(TACT)-z(Treg).

**Fig. 4.**
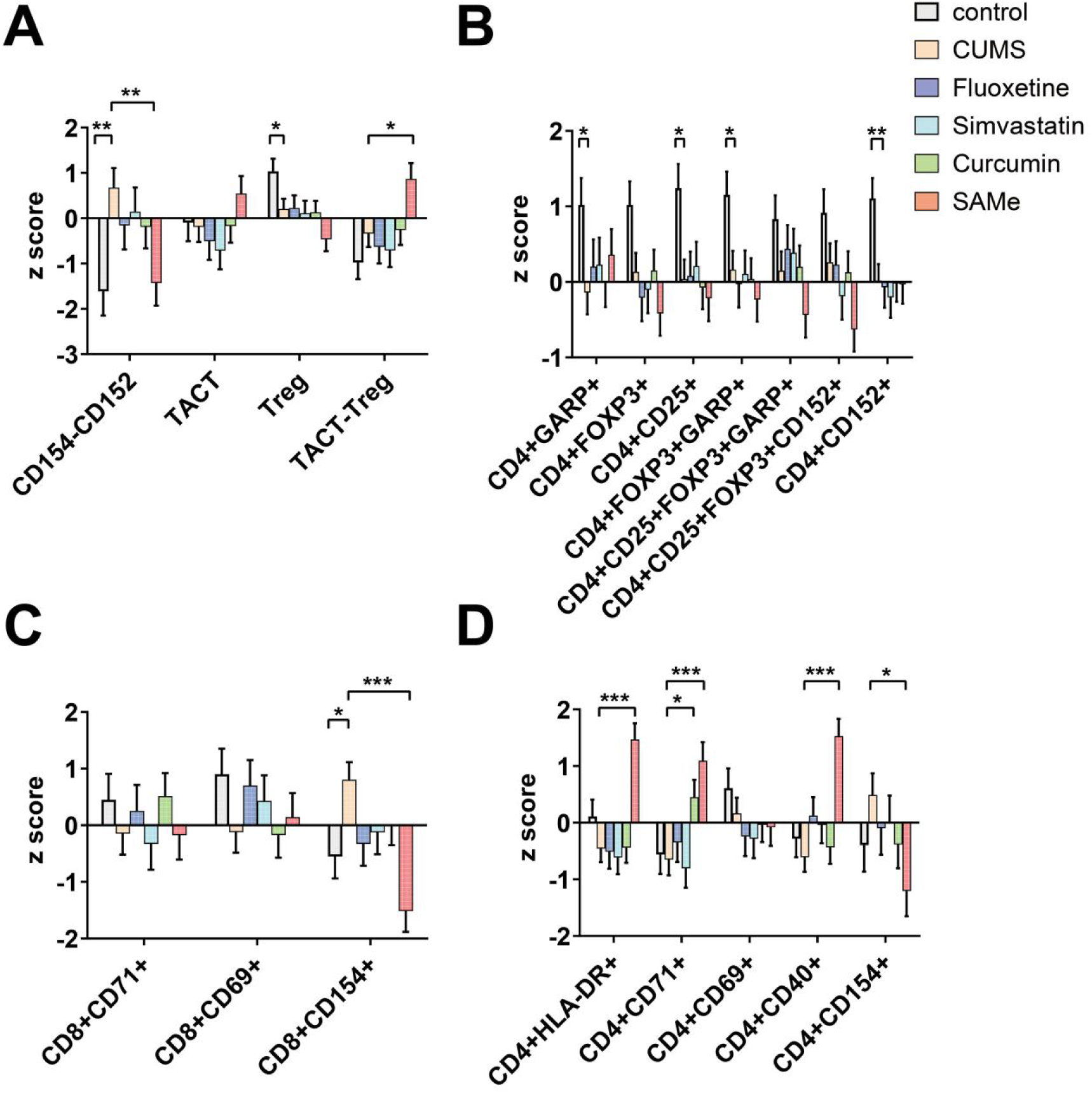
Differences in immune cells among six groups of mice: control, CUMS, fluoxetine, simvastatin, curcumin, and SAMe. The main focus is on the differences in each indicator when comparing CUMS with control, fluoxetine, simvastatin, curcumin, and SAMe. *, *p* < 0.05; **, *p* < 0.01; ***, *p* < 0.001. **(A)** Differences in composite indicators among the six groups. The indices include z(CD154-CD152), which is calculated as z(CD4+CD154+) + z(CD8+CD154+) − z(CD4+CD152+), z(TACT), which represents T-cell activation, calculated as z(CD4+HLA-DR+) + z(CD4+CD71+) + z(CD4+CD69+) + z(CD4+CD40+) + z(CD4+CD154+), and z(Treg), which represents regulatory T cells, calculated as z(CD4+FOXP3+GARP+) + z(CD4+CD25+FOXP3+GARP+) + z(CD4+CD25+FOXP3+CD152+). Furthermore, the ratio of activated T versus Treg cells or z(TACT-Treg) is computed as z(TACT)-z(Treg). **(B)** Differences in regulatory CD4+ cells among the six groups, including CD4+GARP+, CD4+FOXP3+, CD4+CD25+, CD4+FOXP3+GARP+, CD4+CD25+FOXP3+GARP+, CD4+CD25+FOXP3+CD152+, and CD4+CD152+. **(C)** Differences in CD8+ cells among the six groups, including CD8+CD71+, CD8+CD69+, and CD8+CD154+. **(D)** Differences in T-cell activation among the six groups, including CD4+HLA-DR+, CD4+CD71+, CD4+CD69+, CD4+CD40+, and CD4+CD154+. Data are presented as mean ± standard error of the mean (SEM) (n = 8 per group; 4 males and 4 females). Statistical analyses were performed using factorial general linear models (GLMs) with treatment group and sex as fixed factors, including the treatment group × sex interaction. When a significant treatment-group effect was detected, protected least significant difference (LSD) tests were used for pairwise comparisons. The P values shown represent pairwise comparisons between the groups indicated by brackets. All tests were two-tailed.

Compared to the Control group, the CUMS group exhibited significantly higher values in the z(CD154-CD152) index (*p* = 0.006) and lower values in the z(Treg) index (*p* = 0.047), indicating immune imbalances in the CUMS mice. Compared to the CUMS group, the SAMe group had a lower z(CD154-CD152) index (*p* = 0.009) and a higher z(TACT-Treg) index (*p* = 0.025). No significant differences were observed in the T-cell activation composite index between the CUMS group and the other groups.

Regarding regulatory CD4+ cells, we found that the CUMS group had lower percentages of CD4+GARP+ (*p* = 0.032), CD4+CD25+ (*p* = 0.014), CD4+FOXP3+GARP+ (*p* = 0.033), and CD4+CD152+ (*p* = 0.009) cells compared to the Control group, as shown in **Fig. 4B**, indicating impaired immune regulation in CUMS group.

In CD8+ cells, particularly in the percentage of CD8+CD154+ cells, the CUMS group exhibited higher percentages than the Control group (*p* = 0.021), and SAMe significantly reversed the changes in CD8+CD154+ (*p* < 0.001), as depicted in **Fig. 4C**. Among activated T cells, we found no significant changes between the control and CUMS groups. However, the CUMS group had consistently lower percentages of CD4+HLA-DR+, CD4+CD71+, and CD4+CD40+ cells compared to the SAMe group (*p* < 0.001). In contrast, the percentage of CD4+CD154+ cells was higher in the CUMS group compared to the SAMe group (*p* = 0.015). Additionally, the percentage of CD4+CD71+ cells was lower in the CUMS group compared to the Curcumin group (*p* = 0.022), as shown in **Fig. 4D**.

### 3.3 Immune Cell Differences Between Sexes

The differences in immune cell markers between male and female mice are shown in **Fig. 5**. **Fig. 5A** presents the differences between sexes in the composite indices z(CD154-CD152), z(TACT), z(Treg), and z(TACT-Treg). Female mice exhibited higher z(CD154-CD152) values than males (*p* = 0.008). Males showed higher z(TACT) and z(Treg) composite indices than females (all *p* < 0.001). No significant differences were found between the sexes in the composite index of z(TACT-Treg).

**Fig. 5.**
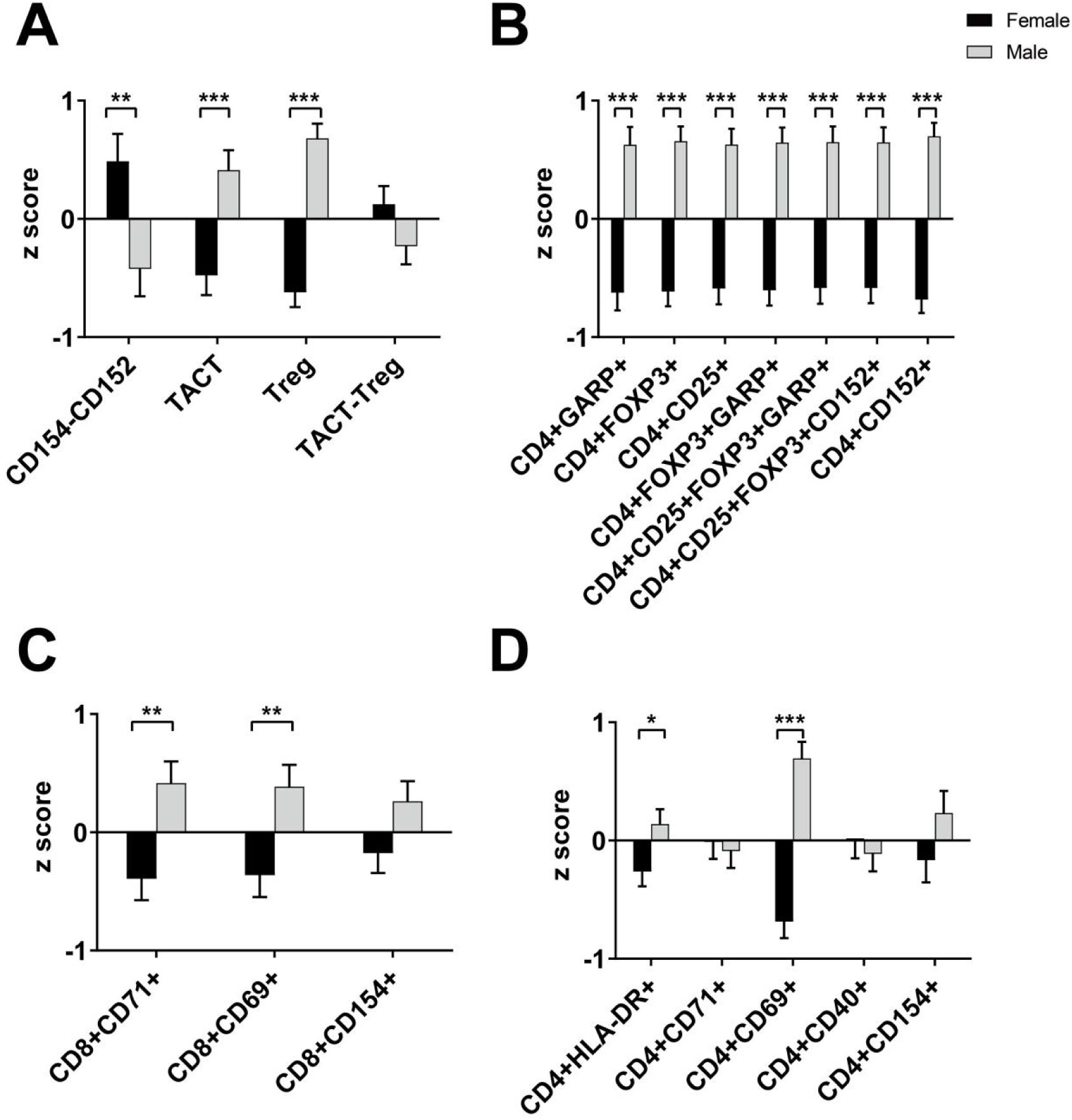
Differences in immune cells between male and female mice. *, *p* < 0.05; **, *p* < 0.01; ***, *p* < 0.001. **(A)** Differences in composite indicators between genders, including z(CD154-CD152), z(TACT), z(Treg), and z(TACT-Treg). A more detailed explanation of the composite indicators can be found in Fig. 4A. **(B)** Differences in regulatory CD4+ cells between genders, including CD4+GARP+, CD4+FOXP3+, CD4+CD25+, CD4+FOXP3+GARP+, CD4+CD25+FOXP3+GARP+, CD4+CD25+FOXP3+CD152+, and CD4+CD152+. **(C)** Differences in CD8+ cells between genders, including CD8+CD71+, CD8+CD69+, and CD8+CD154+. **(D)** Differences in T-cell activation between genders, including CD4+HLA-DR+, CD4+CD71+, CD4+CD69+, CD4+CD40+, and CD4+CD154+. Bars represent mean ± standard error of the mean (SEM) (n = 24 per sex; 4 mice of each sex from each experimental group). Each immune-cell outcome was analyzed using a factorial general linear model (GLM) with treatment group and sex as fixed factors, including the treatment group × sex interaction. The *p* values shown represent the main effect of sex. All tests were two-tailed.

Regarding regulatory CD4+ cells, males consistently showed higher percentages of CD4+GARP+, CD4+FOXP3+, CD4+CD25+, CD4+FOXP3+GARP+, CD4+CD25+FOXP3+GARP+, CD4+CD25+FOXP3+CD152+, and CD4+CD152+ cells than females (*p* < 0.001), as illustrated in **Fig. 5B**.

Among CD8+ T-cell subsets, the proportions of CD8+CD71+ (*p* = 0.003) and CD8+CD69+ (*p* = 0.007) cells were significantly higher in males than in females (**Fig. 5C)**.

In activated T cells, males exhibited higher percentages of CD4+HLA-DR+ (*p* = 0.028) and CD4+CD69+ (*p* < 0.001) cells compared to females, as depicted in **Fig. 5D**. There was no significant sex-by-group (treatment) interaction in any of the immune cell subsets.

## 4. Discussion

This study highlighted the immune dysregulation observed in the CUMS mouse model and its varying responses to pharmacological interventions and sex differences. The CUMS mice exhibited an elevated CD154-CD152 index and a reduced Treg index, indicating a breakdown of immune homeostasis with an imbalance between depleted regulatory CD4+ cells and a relative increase in effector T cells. The administration of fluoxetine, simvastatin, curcumin, and SAMe significantly improved SPT, autonomous activity ability, and anxiety-like behaviors. Fluoxetine and SAMe reduced the immobility time, and SAMe further improved learning and memory impairments. Importantly, the treatments did not normalize the CUMS-induced changes in immune homeostasis. In fact, fluoxetine, curcumin, and simvastatin did not have any significant effects on the CUMS-induced immune subsets, whilst SAMe administration produced distinct immunomodulatory effects. Female mice exhibited lower levels of immunoregulation and indicators of T-cell activation compared with males. Moreover, females showed a significant immune imbalance, as indicated by a higher CD154-CD152 index.

Compared to the control group, CUMS mice showed a significant imbalance between T-cell activation and immune tolerance. This was reflected by an increased CD154-CD152 index (elevated CD8+CD154+ T cells and reduced CD4+CD152+ cells) and a decreased Treg index, consistent with previous research on immune alterations in depression ^1–5^. CD154 (CD40L) is a key T-cell activation marker that promotes immune responses by binding to CD40, while CD152 (CTLA-4) serves as a negative regulator to inhibit immune responses and maintain immune tolerance ^1–6^. The changes in the CD154-CD152 index reflect the imbalance between CD4 and CD8 T cell activation via CD40L and regulation via CTLA-4. Therefore, chronic stress might trigger immune responses through this imbalance, thus contributing to the development of depressive-like symptoms in the CUMS model and major depression in humans ^3,7–10^. Additionally, the reduction of Tregs in CUMS mice reflects breakdown of immune tolerance due to prolonged stress, further exacerbating immune dysregulation. This finding aligns with previous studies, supporting the theory that chronic stress is associated with immune system disturbances in depression ^3,7–10^.

In contrast, the activation status of CD4+ T cells did not show significant effects of CUMS, although CUMS increased the CD154-CD152 index. This may be due to the significant effects of CUMS leading to increases in CD8+CD154+ cell numbers, while depleting CD4+CD152+ T cells. Previous studies have suggested that immune tolerance of CD4+ T cells may be mediated through various pathways, including Treg functions and intracellular signaling ^37^. Moreover, the activation status of CD4+ T cells may also be influenced by external factors, such as antigen specificity and chronic infections, which could lead to variations in immune activation levels ^38^.

Against this background of CUMS-induced immune dysregulation, the study found that fluoxetine, simvastatin, curcumin, and SAMe significantly alleviated anhedonia, reversed reductions in spontaneous activity, and improved OFT-derived anxiety-like behaviors in CUMS mice, with fluoxetine and SAMe reducing the helplessness state and SAMe further improving learning and memory impairments. SAMe, in particular, appeared to improve depressive-like behaviors in association with modulating the CD154 versus CD152 ratio. This was the first study to show that SAMe significantly reduced the CD154-CD152 index by reversing the changes in CD4+CD154+ and CD8+CD154+ cell numbers, thereby restoring immune balance. Previous studies suggest that SAMe might exert its antidepressant effects by restoring neurotransmitter levels (such as serotonin and dopamine) and membrane fluidity ^26^. While no differences were observed in the T-cell activation index and Treg index between the SAMe group and the CUMS group, the TACT-Treg index was elevated following the administration of SAMe. This suggests that SAMe produced mixed immunomodulatory effects, including elevations in activated T cells such as CD4+HLA-DR+, CD4+CD71+, and CD4+CD40+ T cells.

Curcumin and simvastatin improved SPT and OFT outcomes but did not significantly affect FST or AAT measures. Although curcumin is thought to suppress activated immune-inflammatory pathways ^23–25^, it did not significantly affect FST or AAT performance in the present study. Moreover, curcumin increased CD4+CD71+ T-cell numbers and failed to reverse the CUMS-induced increase in the CD154/CD152 ratio and depletion of Tregs, suggesting incomplete immunomodulatory activity under the present experimental conditions.

For simvastatin, a previous mouse CMS study observed more consistent behavioral benefits at 20 mg/kg than at 10 mg/kg, suggesting that dose may partly contribute to the different behavioral responses ^39^.

Antidepressants such as SSRIs (e.g., fluoxetine) may have immunosuppressive activities by attenuating M1 macrophage and T helper-1 activation, and increasing the production of immunoregulatory Treg cytokines (namely IL-10) ^18,19^. Nevertheless, the current study was unable to establish significant effects of fluoxetine on any of the immune subsets. These differences may be attributed to the varying effects of these drugs on peripheral blood mononuclear cells ^17–19^ compared to immune cells in the spleen (the current study). Overall, the differing behavioral and immune responses across treatments are consistent with the NIMETOX framework, in which neuroimmune, metabolic, and oxidative pathways may contribute to treatment effects to varying degrees ^16^.

In addition to these treatment-related patterns, sex further shaped the balance between T-cell activation and immune regulation in CUMS mice. Specifically, this is the first study showing that female mice exhibited a significantly higher CD154-CD152 index, indicating that females may be more prone to breakdown of immune tolerance. In contrast, males exhibited significantly higher levels of Treg index and regulatory CD4+ cells (e.g., CD4+GARP+, CD4+FOXP3+), suggesting that males possess stronger immune regulatory capabilities, consistent with previous research ^28^. Additionally, male mice showed higher levels of the TACT index, activated T cells (such as CD4+HLA-DR+, CD4+CD69+), and CD8+ cells (e.g., CD8+CD71+, CD8+CD69+), reflecting more active immune system activation compared to females, which contradicts some prior findings ^28^.

These sex-linked differences may be due to the immune regulatory effects of sex hormones. Androgens (such as testosterone) are generally thought to exert an immunoregulatory effect on the immune system, thereby maintaining immune tolerance ^30^. However, under chronic stress, androgens might enhance CD8+ T cells and activated T cell proliferation, thus promoting immune responses and maintaining appropriate immune activation levels. This immune adaptive regulation mechanism may enable male mice to maintain effective immune responses during immune challenges by increasing cytotoxic immunity and T-cell activation, leading to higher levels of CD8+ cells and activated T cells ^40–42^. While estrogens generally enhance immune activation, their role under chronic stress may be more complex, potentially both enhancing and suppressing specific immune responses through modulation ^30,43^. Therefore, the role of androgens in immune responses may primarily focus on maintaining immune system balance and moderate activation during immune challenges, making male mice more pronounced in immune activation.

## 5. Conclusions

This study demonstrated that CUMS mice exhibited a breakdown of immune tolerance due to depletion of Tregs and an increase in the CD154/CD152 ratio. Fluoxetine, simvastatin, curcumin, and SAMe partially improved depressive-like behaviors. Nevertheless, the antidepressant-like effects of these four drugs were not mediated by the normalization of CUMS-induced Treg suppression. The antidepressant-like effects of SAMe, reflected by significant improvements in the SPT, FST, OFT, and AAT, may have been partly mediated via normalization of the CD154/CD152 ratio. Moreover, sex differences played an important role in immune responses, with female mice showing a breakdown of immune homeostasis following CUMS. In contrast, male mice showed stronger immune regulatory capabilities and moderate immune activation.

However, this study has some limitations. Although the study observed the effects of different treatments on immune cells in the spleen, the detailed mechanisms underlying specific immune pathways still require further investigation. An additional limitation is the absence of matched vehicle control groups for 1% DMSO and peanut oil. Considering the prior use of these vehicles in published animal studies, separate vehicle-only groups were not included to avoid the use of additional animals, in accordance with the Reduction principle of the 3Rs. Nevertheless, potential vehicle-related effects cannot be completely excluded. Future studies should incorporate histopathological assessments of hippocampal regions and the architecture of the spleen and thymus to complement the behavioral and immunological findings.

## Clinical trial number

Not applicable.

## Ethics approval

All procedures were approved by the Institutional Animal Care and Use Committee (IACUC) of Sichuan Junhui Biotech Co., Ltd. (approval number: IACUC-202411-4-001).

## Consent to participate

Not applicable.

## Consent for publication

Not applicable.

## Acknowledgments

A preprint has previously been published ^44^. We thank the staff of Sichuan Junhui Biotech Co., Ltd for providing the animal facility and Sichuan Youngster Technology Co., Ltd for performing the flow cytometry analysis.

## Author’s contributions

Mengqi Niu: visualization, writing - original draft. Yingqian Zhang: conceptualization, writing - review and editing. Abbas F. Almulla, Tangcong Chen, Yueyang Luo and Mengdie Li: methodology. Juan Li, Yaoyin Zhang and Yulan Huang: Funding acquisition. Michael Maes: conceptualization, formal analysis, writing - review and editing. All authors approved the submitted manuscript.

## Data access statement

The database created during this investigation will be provided by the corresponding author (MM) upon a reasonable request once the authors have thoroughly used the data set.

