## Supplementary Table for "Regulatory T and activated T cells in the chronic unpredictable mild stress mice: sex differences and effects of treatments"

**Electronic Supplementary File (ESF)**

**ESF Table 1**. Pharmacological interventions: formulations, dosing, and administration schedule.

| **Group** | **Compound** | **Supplier** | **CAS No.** | **Dose** | **Route** | **Vehicle / formulation** | **Treatment window** | **Citation** |
| --- | --- | --- | --- | --- | --- | --- | --- | --- |
| Control + Saline | Vehicle control | — | — | — | Oral gavage | 0.9% Saline | q.d., weeks 3–6 (28 d) | — |
| CUMS + Saline | Vehicle control | — | — | — | Oral gavage | 0.9% Saline | q.d., weeks 3–6 (28 d) | — |
| CUMS + Fluoxetine | Fluoxetine | Aladdin (Shanghai, China) | 54910-89-3 | 20 mg/kg | Oral gavage | 0.9% Saline | q.d., weeks 3–6 (28 d) | Li et al., 2022 (1) |
| CUMS + Simvastatin | Simvastatin | Aladdin (Shanghai, China) | 79902-63-9 | 10 mg/kg | Oral gavage | 1% DMSO | q.d., weeks 3–6 (28 d) | Yan et al., 2020 (2); Adeli et al., 2019 (3) |
| CUMS + Curcumin | Curcumin | Aladdin (Shanghai, China) | 458-37-7 | 20 mg/kg | i.p. | Peanut oil (suspension) | q.d., weeks 3–6 (28 d) | Kulkarni et al., 2008 (4); Carroll et al., 2010 (5) |
| CUMS + SAMe | S-adenosylmethionine (SAMe) | Solarbio (Beijing, China) | 97540-22-2 | 100 mg/kg | Oral gavage | 0.9% Saline | q.d., weeks 3–6 (28 d) | Beauchamp et al., 2020 (6) |

Starting 2 weeks after CUMS initiation (experimental week 3), the CUMS-exposed mice were randomized into five groups (n = 12/group, sex-balanced), while the non-stressed control group remained as a separate sixth group. All six groups received the corresponding drug or vehicle once daily for 28 days (weeks 3–6). All drug solutions/suspensions were freshly prepared under dim light.

Abbreviations: CUMS, chronic unpredictable mild stress; SAMe, S-adenosylmethionine; i.p., intraperitoneal; DMSO, dimethyl sulfoxide; q.d., once daily.
